# *De novo* Design of Peptides that Bind Specific Conformers of α-Synuclein

**DOI:** 10.1101/2023.11.14.567090

**Authors:** Hailey M. Wallace, Hyunjun Yang, Sophia Tan, Henry S. Pan, Rose Yang, Junyi Xu, Hyunil Jo, Carlo Condello, Nicholas F. Polizzi, William F. DeGrado

**Author notes:** **Corresponding Author William F. DeGrado**, Department of Pharmaceutical Chemistry and the Cardiovascular Research Institution, University of California, San Francisco, San Francisco, CA 94158, USA.

## Abstract

Insoluble amyloids rich in cross-β fibrils are observed in a number of neurodegenerative diseases. Depending on the clinicopathology, the amyloids can adopt distinct supramolecular assemblies, termed conformational strains. However, rapid methods to study amyloid in a conformationally specific manner are lacking. We introduce a novel computational method for *de novo* design of peptides that tile the surface of α-synuclein fibrils in a conformationally specific manner. Our method begins by identifying surfaces that are unique to the conformational strain of interest, which becomes a “target backbone” for the design of a peptide binder. Next, we interrogate structures in the PDB database with high geometric complementarity to the target. Then, we identify secondary structural motifs that interact with this target backbone in a favorable, highly occurring geometry. This method produces monomeric helical motifs with a favorable geometry for interaction with the strands of the underlying amyloid. Each motif is then symmetrically replicated to form a monolayer that tiles the amyloid surface. Finally, amino acid sequences of the peptide binders are computed to provide a sequence with high geometric and physicochemical complementarity to the target amyloid. This method was applied to a conformational strain of α-synuclein fibrils, resulting in a peptide with high specificity for the target relative to other amyloids formed by α-synuclein, tau, or Aβ40. This designed peptide also markedly slowed the formation of α-synuclein amyloids. Overall, this method offers a new tool for examining conformational strains of amyloid proteins.

## Introduction

Insoluble amyloid aggregates are hallmarks in the pathogenesis of neurodegenerative and systemic amyloidosis diseases.^1^ The formation of amyloid fibrils occurs through the self-assembly of amyloidogenic proteins, namely, amyloid-β (Aβ), tau, and α-synuclein (αSyn). These amyloidogenic proteins aggregate after aberrant conformational changes and misfolding, resulting in an accumulation of β-strand-rich, insoluble fibrils.^2,3^ The misfolded monomers in the fibrils stack against one another and interdigitate as β-sheets, forming a cross-β supramolecular motif.^4^ These amyloid structures have been linked to a diverse range of disorders, making them a focal point of scientific investigation. For example, αSyn amyloids accumulate as inclusion bodies in the neurons of Parkinson’s disease patients ^5, 6^, which are correlated with cellular dysfunction, neuronal damage, and neuroinflammation, leading to the progressive loss of cognitive and motor functions.^7,8^

With the advancement of solid-state nuclear magnetic resonance (ssNMR) and cryo-electron microscopy (cryo-EM), the structures of many fibrils at the molecular resolution are now available. ^9,10,11.12,^ Depending on clinicopathology, each amyloidogenic protein can adopt multiple, distinct supramolecular assemblies, termed conformational strains.^13,14,15^ This conformational strain polymorphism of each amyloidogenic protein adds complexity to study amyloid-related disorders and, in some cases, multiple conformational strains are associated with a single disease (e.g. two conformers of tau amyloids are observed in Alzheimer’s disease patients^16^). A method that could enable the identification and association of specific conformational strains at low cost (relative to, e.g., cryo-electron microscopy) would provide valuable insight into the nature of amyloidopathies. The development of amyloid conformational-specific binders will advance our fundamental understanding of the molecular recognition of amyloid structure for therapeutic strategies. Such binders will also enable the identification of polymorphs *in vitro* and in animal models, providing valuable reagents for histological examination of brain tissue. Ultimately, the knowledge gained in this process can contribute to the design of molecular therapeutics for the recognition and/or degradation of fibrils in a cellular context.

Current approaches have used small molecules, peptides, and antibodies as tools to recognize fibrils and disrupt their aggregation kinetics. ^17,18,19,20,21,22,23,24,25,26^ However, these approaches exhibit varying degrees of conformational specificity. There are also molecular dyes that recognize differences in conformational strains and shift excitation/emission profiles; however, they often bind to multiple types of fibrils.^27,28,29^ Progress has been made in the field of protein design to engineer high-affinity fibril capping peptides that inhibit amyloid growth.^23,24,25,26^ However, these capping sites are limited in a growing fibril when compared to the immense fibril surface, providing only limited surface area for specific molecular recognition. These limitations underscore the feasibility of de novo designed binders targeting the surface of fibrils.

In this work, we have developed a computational method to design peptides that bind amyloids in a conformational strain-specific manner. We focused on *in vitro* fibrils for this model study because they are available in quantities for bulk studies and do not require biosafety requirements associated with handling human tissues. The designed peptides interact in a structure-specific manner with amyloids, and amyloid growth. Finally, our new computational approach should, in principle, allow for the development of new tools for the structure-based design of proteins that bind to other repeating protein surfaces.

## Results

### Computational Design

We began by identifying possible discriminating binding epitopes along the fibril surface of αSyn. Cryo-EM structures of *in vitro*-prepared WT^30^ and E46K^31^ αSyn fibrils produced under the same conditions revealed two distinct conformational strains (**Figure 1**). For generality, we sought regions that were identical in sequence between these two amyloids but differed in their conformation and solvent exposure of amino acid side chains. To facilitate the identification of regions that vary considerably between the two strains, we computed the solvent accessible surface areas (SASA) for each residue of αSyn in both conformations.^32^ By aligning the SASA scores based on the sequence, we identified a distinguishable region spanning residues 62–66 that shared a identical sequence but had markedly different exposures to solvent between the two conformational strains. In particular, Gln62, Thr64, and Val66 are solvent-exposed in the WT αSyn structure, while in the mutant E46K αSyn, they are buried in the fibril core (**Figure 1**). Although the mutation occurs at residue 46 in E46K, this residue is rather buried, ensuring that any discrimination we observe is not due to this residue but instead reflects conformational differences in the rest of the protein.

**Figure 1.**
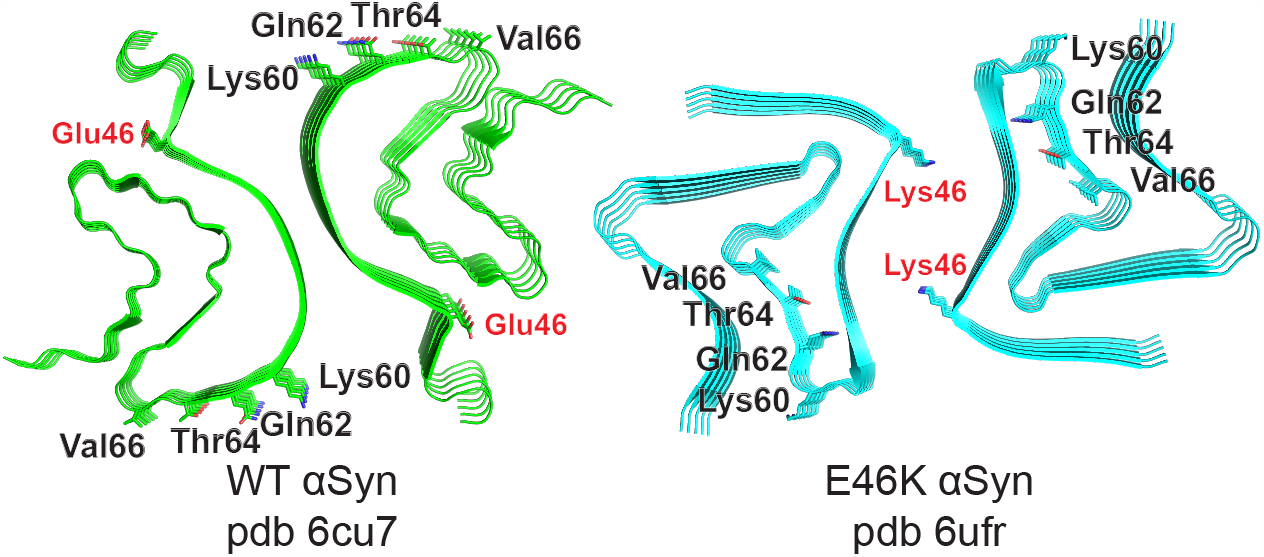
Cryo-EM structures of WT (green) and E46K (cyan) αSyn fibrils. The most discriminating residues Gln62, Thr64, and Val66 project outward in 6cu7 but inward in 6ufr. E46 and K46 are buried in each structure.

To construct the target binding surface, we stacked the 5-residue fragment of the distinguishable region along the Z-axis, using the screw symmetry of the fibril cryo-EM structure (**Figure 2**). We then extracted three neighboring strands and used the program MASTER^33^ to search for structures in the protein structural databank (PBD) that contained similar backbone coordinates to this motif. Because 3-stacked β-strands are highly prevalent in structures in the PDB, a large number of matches were retrieved (15395 matches at 1.0 Å rmsd). We then extracted the secondary structural motifs that are in the vicinity of the 3-strand motif (i.e., the target binding surface). In this process, interacting β-sheet motifs were filtered out because we expected that any hits conforming to the β-sheet conformation of an amyloid would probably self-assemble into an amyloid, even in the absence of αSyn fibrils. This resulted in a large number of helical poses, which were clustered based on the geometry of the interacting helix, its orientation relative to the target binding surface, and its frequency of occurrence (designability).

**Figure 2.**
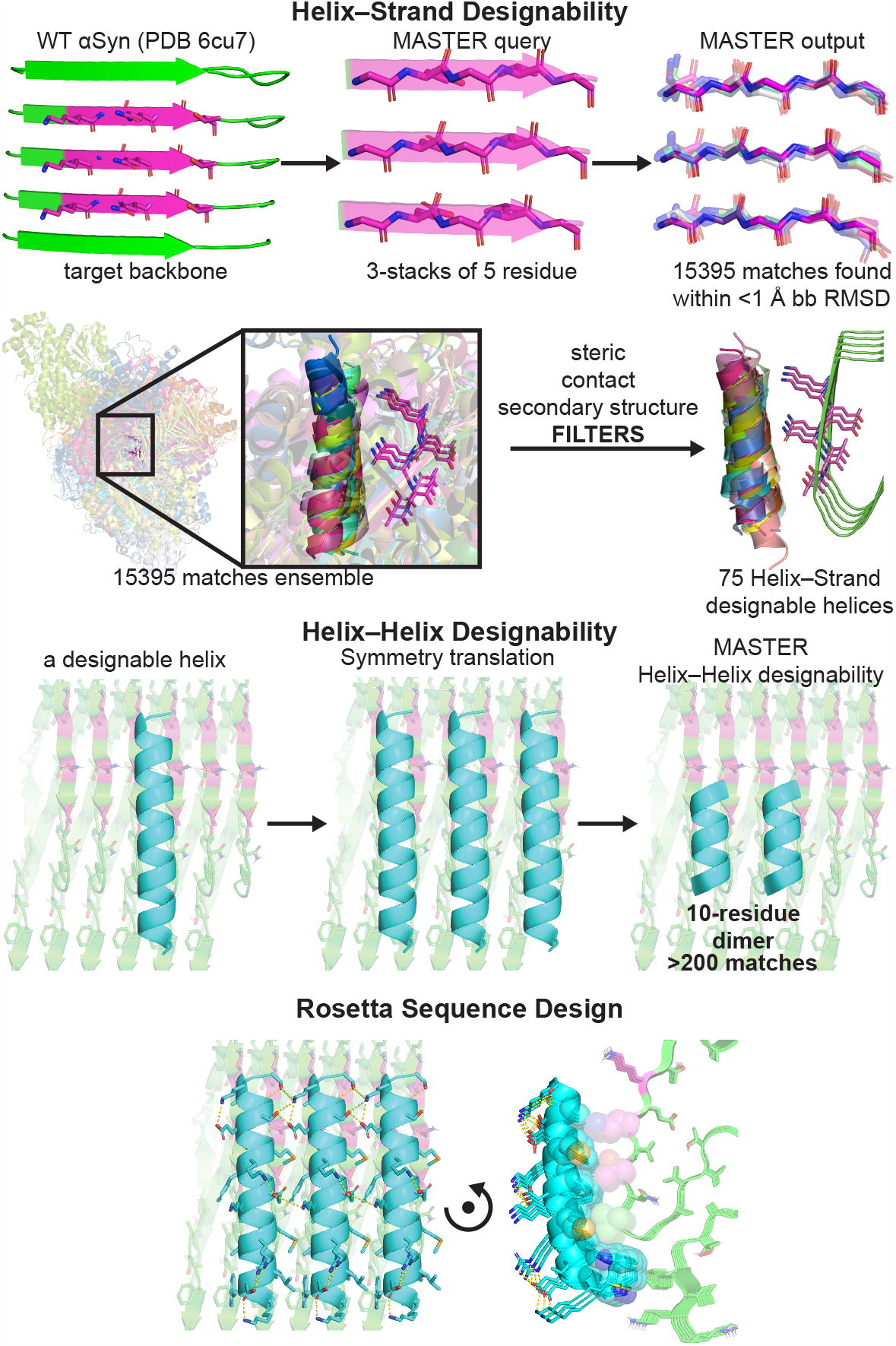
Amyloid peptide binder design pipeline. (A) Helix–strand designability is evaluated using MASTER and structural filters consisting of steric, contact, and secondary structure. This produced helix backbones favorable for helix–strand interaction. (B) Helix–helix designability is evaluated by applying the screw symmetry of amyloid structure on the helix and assessing their occurrence in the PDB database using MASTER. (C) The peptide sequence was generated by Rosetta using the symmetry files, considering both interactions of helix–strand and helix–helix while fixing the amyloid residues.

Having identified helical poses that could interact favorably with the underlying amyloid surface, we next used the screw symmetry of the fibril to create the overall assembly and design helix–helix interactions. The repeating geometry of the underlying fibrils dictated the stoichiometry and topology of the peptides relative to the target amyloid surface. The spacing of the β-strands is approximately 5 Å, restrained by the formation of inter-strand mainchain hydrogen bonds. On the other hand, the interhelical distance of helices within water-soluble proteins is approximately 8–11 Å, depending on the bulk of the residues lining the interface.^34^ These values dictated a stoichiometry of one helix for every two β-strands. Application of the fibril symmetry created a helical array with 9.6 Å spacing (twice the 4.8 Å spacing of the β-strands in the fibril). Moreover, the symmetry dictated a parallel orientation between the interacting helices.

Parallel packing of helices with a 10 Å spacing occurs quite frequently in proteins^34^, suggesting that this should be a highly designable interface and may facilitate cooperative peptide binding on the amyloid surface. However, other interhelical parameters such as the α-helical phase and the precise interhelical crossing angle should also be favorable in the design.^35^ Thus, to identify helices suitable for helix–helix stacking along the amyloid surface, we again used MASTER to query two neighboring helices. The number of matches determined the feasibility of each helix–helix geometry, resulting in 75 helical backbones (>200 matches at 1.0 Å rmsd) that were well suited for the joint optimization of helix–strand and helix–helix interfaces in the next step of design.

We then ran a flexible backbone sequence design protocol using symmetry files for symmetric helix–helix packing within Rosetta.^36,37,38^ Rosetta symmetry files enable sequence design considering the symmetry of the fibril and the peptides, engendering favorable helix–strand and helix–helix interactions as a joint optimization problem. A total of 75,000 designs were generated using flexible backbone design. To rank the designs, we used Rosetta’s REF2015 score function. We took the energy score of [six β-strands • one helix] and subtracted the sum of [six β-strands] and [one helix] to estimate the helix–strand interaction. The helix–helix interaction score was similarly computed as the contribution from interactions in a symmetrical complex (**Figure S1**). During the selection process, designs were further filtered to minimize unsatisfied hydrogen bonds, ensuring only the most promising candidates were retained.

### Experimental evaluation

The top four designs derived from three helical motifs were selected for synthesis and experimental evaluation (**Table 1**). Additionally, we synthesized a scrambled control peptide, in which the sequence of peptide 4 was randomized. Each peptide was synthesized using the conventional Fmoc-solid phase peptide synthesis method with and without the *N*-terminus capped with fluorescein-5-isothiocyanate (FITC). A β-Ala was included as a linker between the peptides and the FITC group to reduce *N*-terminal cleavage by FITC.^39^ All synthesized peptides showed good solubility in aqueous solution.

**Table 1.**
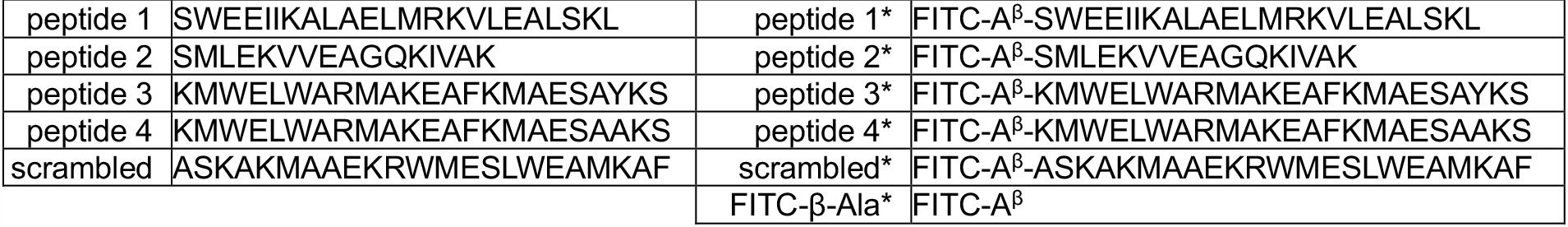
Sequences of designed and synthesized peptides.

We examined the binding of our peptides to WT αSyn fibrils using confocal microscopy. Peptides 1*–4* were dissolved and diluted in PBS buffer at concentrations ranging from 10 nM to 0.1 nM with a final volume of 50 μL in a 384-well plate (**Figure 3A**). To each well, 1-μL aliquot of 300 μM WT αSyn fibril was added and incubated for 30 min. The plate was centrifuged to pellet the fibrils for confocal microscopy. At 10 nM concentration, Peptide 4* displayed strong intensity indicating good binding. Even at concentrations as low as 1 nM, binding was still detectable with a good signal-to-noise ratio. The other three peptides 1*–3*, however, did not stain the fibrils brightly. It is interesting to note that peptide 3* was not bright despite having only a single alanine to tyrosine mutation from peptide 4*. Therefore, we focused additional characterization on peptide 4*.

**Figure 3.**
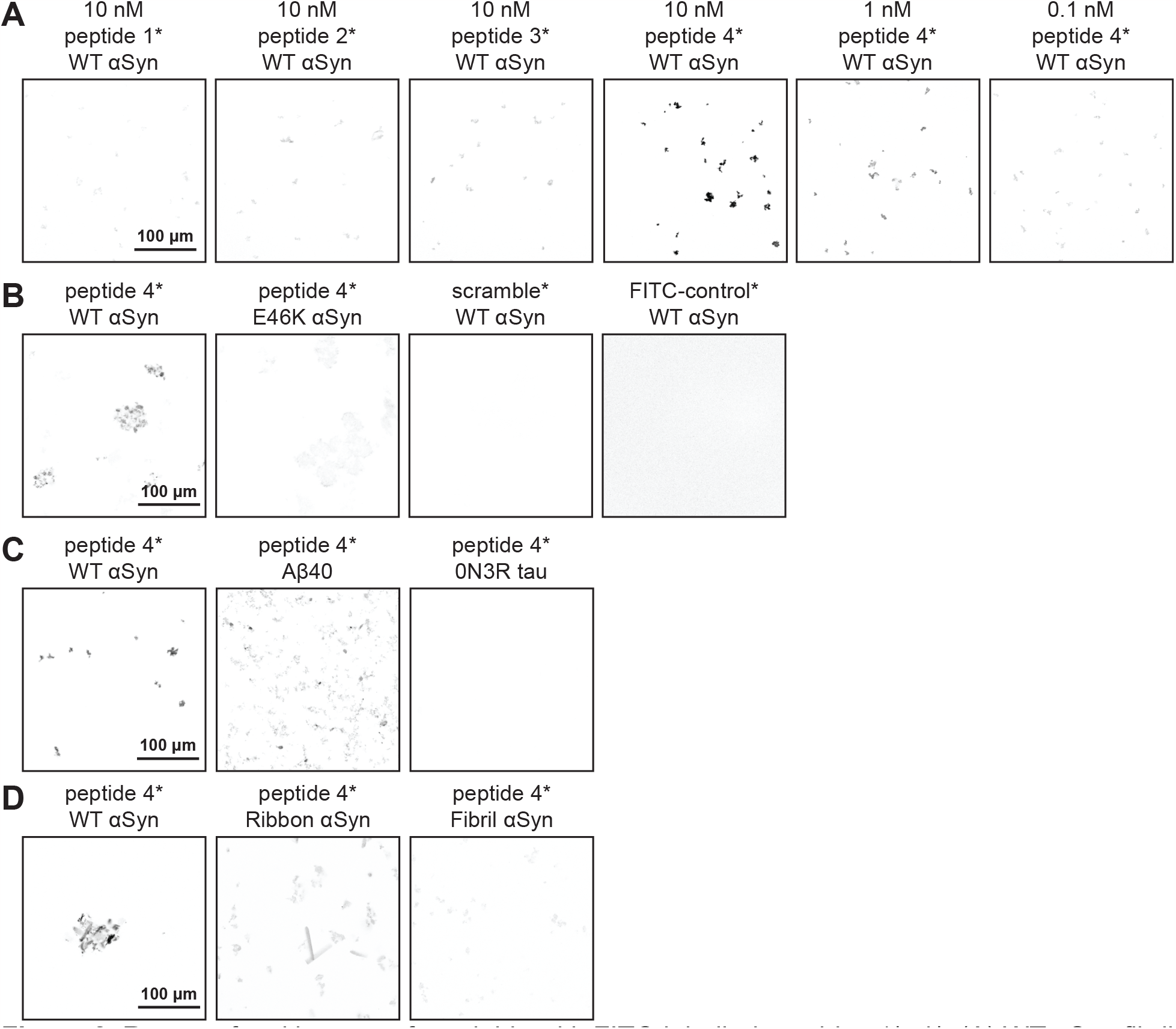
Raw confocal images of amyloids with FITC-labelled peptides 1*–4*. (A) WT αSyn fibril with peptides 1*–4*. (B) Peptide 4* with WT αSyn and E46K αSyn fibrils. (C) Peptide 4* with WT αSyn, Aβ40 and 0N3R tau fibrils. (D) Peptide 4* with three conformers of αSyn fibrils.

To confirm the specificity of peptide 4* binding to WT αSyn fibrils, we compared the fluorescence intensity with E46K αSyn (**Figure 3B**). Since the targeting region comprised of Gln62, Thr64 and Val66 is buried in the E46K αSyn fibril structure, no bright signal was expected. Indeed, we observed bright signal from the WT αSyn fibrils whereas only weak fluorescence from E46K αSyn which matched the intensity from the scrambled* peptide. These data suggested that peptide 4* bound WT αSyn fibrils with good affinity and did not bind E46K αSyn fibrils validating our peptide design method to bind structure-defined amyloid conformer of interest. To rule out any nonspecific binding of the FITC dye, we also synthesized FITC-β-Ala, and no signal was observed with αSyn fibrils. We then examined if binding occurred with other types of amyloids such as Aβ40 and 0N3R tau fibrils (**Figure 3C**). Peptide 4* showed dim fluorescence for Aβ40 and a low signal for 0N3R tau fibrils, in comparison to bright WT αSyn. Collectively, these data suggest that peptide 4* recognizes the targeted WT αSyn fibril conformational strain over E46K as well as other amyloid fibrils.

We then investigated whether peptide 4* could distinguish conformational strains of αSyn based solely on the structural differences, where the conformational strains each share identifical amino-acid sequences. Three different wildtype αSyn fibril conformers (named WT, Ribbon, and Fibril) are produced in three different buffer conditions.^40^ Peptide 4* showed good intensity for the targeted WT αSyn conformer, dim signal for αSyn Ribbon, and low signal for αSyn Fibril, respectively (**Figure 3D**). These data clearly demonstrate that peptide 4* can effectively distinguish structural differences between conformational strains of the same amyloid sequence, highlighting the power of our peptide-design pipeline.

To gain structural insight into the peptide-amyloid complex, we first turned to protein– protein docking protocols from MOE and Hdock^41^ to predict the potential binding poses of the peptide bound to the target amyloid structure (**Figure 4A**). When three copies of the designed helix were rigidly docked onto six β-strands of the fibril, the two methods were in good agreement with the design. A backbone rmsd difference of only 0.8 Å (relative to the computational design) was observed for peptide 4 when the helices were docked using Hdock and only slightly larger using MOE.

**Figure 4.**
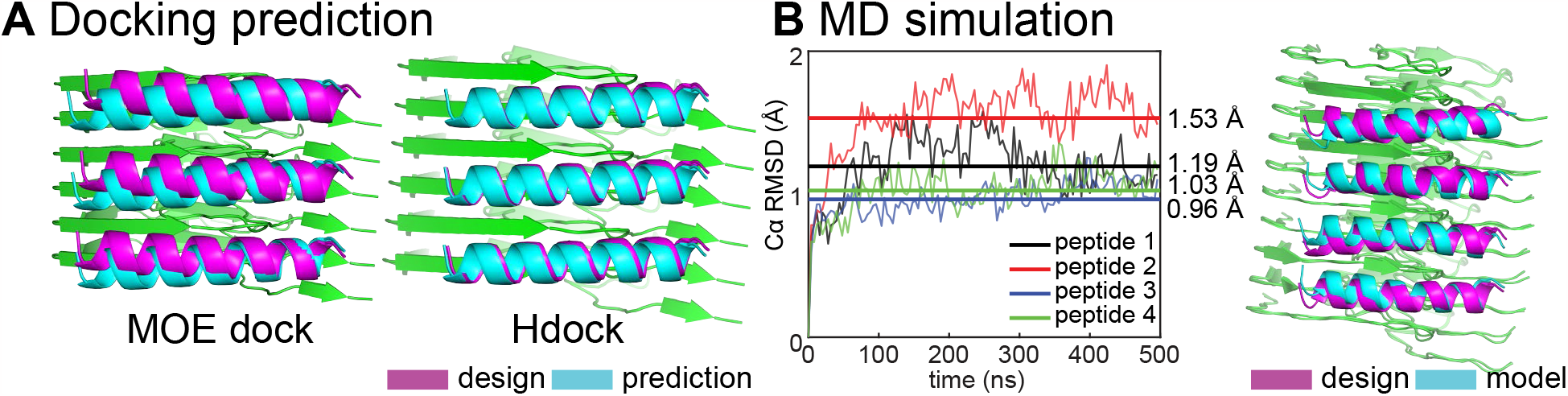
(A) Docking prediction of the designs using MOE dock and Hdock. The backbone rmsd for design vs prediction from MOE dock was 2.0 Å and 0.8 Å for Hdock. (B) MD simulation of design and the WT αSyn fibril (PDB 6cu7) with a plot showing the Cα rmsd calculation over 500 ns simulation.

We next used 500 ns molecular dynamics (MD) simulations to assess the stability of the designed complex (**Figure 4B**). Four copies of designed helices in an array were simulated in complex with ten β-strand chains for 500 ns and their helical backbone rmsds were calculated to monitor stability of the initial configuration on this time scale. Peptides 1 and 2, which did not show appreciable binding in the confocal microscopy study, showed the largest deviations from the initial model. The functional peptide 4 was quite stable by this metric, indicating that MD was able to differentiate this binder from non-functional peptides. However, the Cα RMSD of peptide 3 was similar to peptide 4, indicating that stability on the 500 nsec time scale was a necessary but not sufficient metric predictor of binding.

To determine whether the binding interaction was stoichiometric, we determined the binding affinity between peptide 4 and WT αSyn fibril using a ligand depletion sedimentation assay (**Figure 5A**). WT αSyn fibrils were titrated into a fixed concentration of peptide 4*, and the amount of peptide remaining in solution was determined after ultracentrifugation. Our results displayed a saturation binding curve, indicating a plateau in binding. The data fits well to a binding isotherm with a KD = 1.3 µM and a stoichiometry of 1 peptide per 2 αSyn monomer, which matched the designed stoichiometry. It is, however, still possible that the peptide initially binds with higher affinity at low peptide/surface ratios and becomes weaker as saturation is approached. In a control experiment with the scramble* peptide, no binding to WT αSyn fibril was observed.

**Figure 5.**
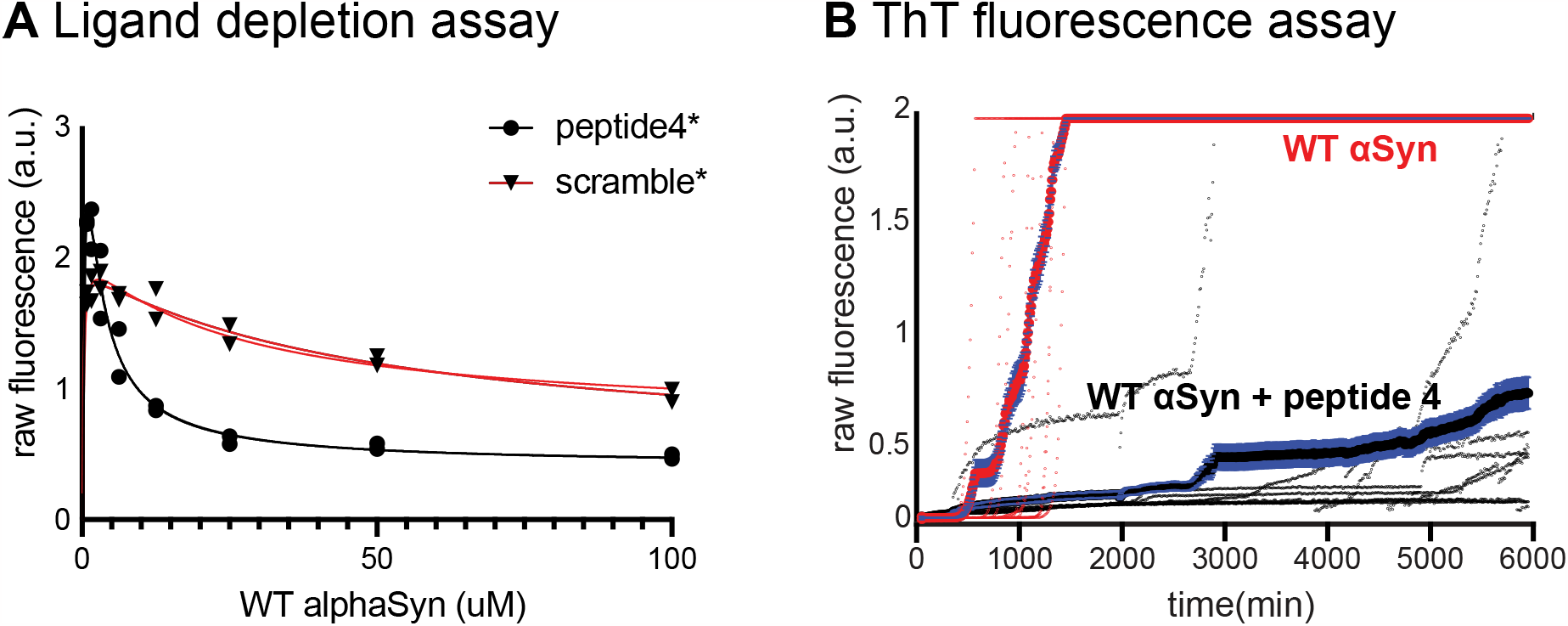
(A) Ligand depletion assay with peptide 4 reveals a saturation binding curve with KD=1.3 μM and binding stoichiometry of 1 peptide per 2 αSyn proteins (black). The scramble* control did not show binding (red) (B) ThT fluorescence assay with 300 μM αSyn monomer with or without 200 μM peptide 4. The mean fluorescence signal from octuplicate is shown in thicker line with standard deviation. A rapid increase in ThT fluorescence is observed at 500 min for αSyn alone whereas a slower increase is observed with the αSyn with peptide 4.

We next assessed the effect of peptide 4 on the rate of fibrillization of αSyn in the presence of thioflavin T (ThT), a fluorogenic probe for amyloid formation (**Figure 5B**). We prepared fibrilization kinetics experiment with WT αSyn monomer in the presence and absence of peptide 4, each in of eight replicates. We noticed an increase in the fluorescence signal for WT αSyn at around 500 min, indicating αSyn fibrillization. This was compared to αSyn fibrilization with peptide 4 where a delayed increase in fluorescence is observed at around 2500 min and a slower increase of fluorescence signal. These data suggest that peptide 4 can interfere with the fibrilization kinetics. We suspect that peptide 4 tiles the solvent exposed fibril surface and prevents the secondary nucleation event.^42^ It is also possible that peptide 4 competes with ThT for the amyloid binding site, or peptide 4 favored one conformational strain over the other polymorphs. Collectively, peptide 4 slowed the fibrilization kinetics and did not act as an inducer.

## Discussion

Our peptide-design approach provides a novel method to identify different conformational strains of amyloid fibrils. For example, αSyn has distinct conformational strains associated with Parkinson’s, multiple system atrophy, and dementia with Lewy bodies. Similarly, one or more distinct conformational strains are specific for tau fibrils in Alzheimer’s vs Pick’s disease or Aβ fibrils in sporadic vs familial Alzheimer’s diseases. Current conformational strain identification methods typically involve a combination of antibodies and dye-staining techniques. The present method utilizes small, robust peptides that can be chemically synthesized. Alternatively, these peptides can be genetically fused to fluorescent proteins for studying the mechanisms of intracellular amyloid formation. They may also offer attractive possibilities for targeted degradation of amyloids.

More generally, this work presents a comprehensive approach for designing peptide or protein assemblies along any structured, repeating biological surface. In this method, the specificity of interaction is generated through both peptide–strand interactions and peptide– peptide interactions. The interaction of a single peptide is expected to be relatively weak, but by forming highly favorable peptide–peptide repeating interactions, the overall free energy of assembly becomes significantly favorable. Moreover, the specific interaction that each adjacent peptide forms with its neighbors stabilizes the geometry required for peptide-strand interactions. Altogether, a highly specific assembly is created, and might be widely useful as such.

## ASSOCIATED CONTENT

### Supporting Information

The Supporting Information is available free of charge on the ACS Publications website at DOI: XXXX.

Procedures for the design pipeline; preparation of αSyn fibrils (WT, E46K, Rod, and Fibril), and Aβ40 and 0N3R tau fibrils; peptide synthesis; ligand depletion sedimentation assay; ThT fluorescence assays; confocal imaging; characterization data (MALDI-MS, CD) for peptide 4.

## AUTHOR INFORMATION

### Author Contributions

H. M. W. and H. Y. contributed equally.

### Notes

The authors declare no competing financial interest.

## ACKNOWLEDGEMENTS

We thank the National Institute of Health (GM122603) for financial support. N.F.P. acknowledges support for NIH (5R00GM135519). H.Y. received postdoctoral fellowship funding from the BrightFocus Foundation (A2020039F). We thank Drs. Benjamin Keith Keitz, David W. Taylor, Evan Schwartz, and Axel Brilot for their contribution in enabling and collecting preliminary data at the Sauer Structural Biology Laboratory. We also thank Drs. Greg Merz and David Boyer for their intellectual contribution on the data analysis.

## REFERENCES

1 Chiti, F.; Dobson, C. M. Protein misfolding, functional amyloid, and human disease. Annu. Rev. Biochem. 2006, 75, 333–366.

2 Nelson, R.; Sawaya, M. R.; Balbirnie, M.; Madsen, A. Ø.; Riekel, C.; Grothe, R.; Eisenberg, D. Structure of the Cross-Beta Spine of Amyloid-like Fibrils. Nature 2005, 435, 773–778.

3 Fitzpatrick, A. W. P.; Debelouchina, G. T.; Bayro, M. J.; Clare, D. K.; Caporini, M. A.; Bajaj, V. S.; Jaroniec, C. P.; Wang, L.; Ladizhansky, V.; Müller, S. A.; MacPhee, C. E.; Waudby, C. A.; Mott, H. R.; De Simone, A.; Knowles, T. P. J.; Saibil, H. R.; Vendruscolo, M.; Orlova, E. V.; Griffin, R. G.; Dobson, C. M. Atomic Structure and Hierarchical Assembly of a Cross-β Amyloid Fibril. Proc. Natl. Acad. Sci. 2013, 110, 5468–5473.

4 Tycko, R. Progress towards a molecular-level structural understanding of amyloid fibrils. Curr. Opin. Struct. Biol. 2004, 14, 96–103.

5 Spillantini, M. G.; Crowther, R. A.; Jakes, R.; Hasegawa, M.; Goedert, M. Alpha-Synuclein in Filamentous Inclusions of Lewy Bodies from Parkinson’s Disease and Dementia with Lewy Bodies. Proc. Natl. Acad. Sci. U. S. A. 1998, 95, 6469–6473.

6 Stefanis, L. α-Synuclein in Parkinson’s Disease. Cold Spring Harb. Perspect. Med. 2012, 2, a009399.

7 Eisenberg, D.; Jucker, M. The Amyloid State of Proteins in Human Diseases. Cell 2012, 148, 1188–1203.

8 Ke, P. C.; Zhou, R.; Serpell, L. C.; Riek, R.; Knowles, T. P. J.; Lashuel, H. A.; Gazit, E.; Hamley, W.; Davis, T. P.; Fändrich, M.; Otzen, D. E.; Chapman, M. R.; Dobson, C. M.; Eisenberg, D. S.; Mezzenga, R. Half a Century of Amyloids: Past, Present and Future. Chem. Soc. Rev. 2020, 49, 5473–5509.

9 Tycko. R. Solid-state NMR studies of amyloid fibril structure. Annu. Rev. Phys. Chem. 2011, 62, 279–299.

10 Tuttle, M. D.; Comellas, G.; Nieuwkoop, A. J.; Covell, D. J.; Berthold, D. A.; Kloepper, K. D.; Courtney, J. M.; Kim, J. K.; Barclay, A. M.; Kendall, A.; Wan, W.; Stubbs, G.; Schwieters, C. D.; Lee, V. M. Y.; George, J. M.; Rienstra, C. M. Solid-State NMR Structure of a Pathogenic Fibril of Full-Length Human α-Synuclein. Nat. Struct. Mol. Biol. 2016, 23, 409–415.

11 Scheres, S. H.; Zhang, W.; Falcon, B.; Goedert, M. Cryo-EM structures of tau filaments. Curr. Opin. Struct. Biol. 2020, 64 17–25.

12 Fitzpatrick, A.W.; Saibil, H. R. Cryo-EM of amyloid fibrils and cellular aggregates. Curr. Opin. Struct. Biol. 2019, 58, 34–42.

13 Cohen, M. L.; Kim, C.; Haldiman, T.; ElHag, M.; Mehndiratta, P.; Pichet, T.; Lissemore, F.; Shea, M.; Cohen, Y.; Chen, W.; Blevins, J.; Appleby, B. S.; Surewicz, K.; Surewicz, W. K.; Sajatovic, M.; Tatsuoka, C.; Zhang, S.; Mayo, P.; Butkiewicz, M.; Haines, J. L.; Lerner, A. J.; Safar, J. G. Rapidly progressive Alzheimer’s disease features distinct structures of amyloid-β. Brain 2015, 138, 1009–1022.

14 Liu, H.; Kim, C.; Haldiman, T.; Sigurdson, C. J.; Nyström, S.; Nilsson, K. P. R.; Cohen, M. L.; Wisniewski, T.; Hammarström, P.; Safar, J. G. Distinct conformers of amyloid beta accumulate in the neocortex of patients with rapidly progressive Alzheimer’s disease. J. Biol. Chem. 2021, 297, 101267.

15 Lau, A.; So, R. W. L.; Lau, H. H. C.; Sang, J. C.; Ruiz-Riquelme, A.; Fleck, S. C.; Stuart, E.; Menon, S.; Visanji, N. P.; Meisl, G.; Faidi, R.; Marano, M. M.; Schmitt-Ulms, C.; Wang, Z.; Fraser, P. E.; Tandon, A.; Hyman, B. T.; Wille, H.; Ingelsson, M.; Klenerman, D.; Watts, J. C. α-Synuclein strains target distinct brain regions and cell types. Nat. Neurosci. 2020, 23, 21–31.

16 Fitzpatrick, A. W. P.; Falcon, B.; He, S.; Murzin, A. G.; Murshudov, G.; Garringer, H. J.; Crowther, R. A.; Ghetti, B.; Goedert, M.; Scheres, S. H. W. Cryo-EM structures of tau filaments from Alzheimer’s disease. Nature 2017, 547, 185–190.

17 Sievers, S. A.; Karanicolas, J.; Chang, H. W.; Zhao, A.; Jiang, L.; Zirafi, O.; Stevens, J. T.; Münch, J.; Baker, D.; Eisenberg, D. Structure-Based Design of Non-Natural Amino-Acid Inhibitors of Amyloid Fibril Formation. Nature 2011, 475, 96–100.

18 Ladiwala, A. R. A.; Bhattacharya, M.; Perchiacca, J. M.; Cao, P.; Raleigh, D. P.; Abedini, A.; Schmidt, A. M.; Varkey, J.; Langen, R.; Tessier, P. M. Rational Design of Potent Domain Antibody Inhibitors of Amyloid Fibril Assembly. Proc. Natl. Acad. Sci. U. S. A. 2012, 109, 19965–19970.

19 Cheng, P.-N.; Liu, C.; Zhao, M.; Eisenberg, D.; Nowick, J. S. Amyloid β-Sheet Mimics that Antagonize Protein Aggregation and Reduce Amyloid Toxicity. Nat. Chem. 2012, 4, 927–933.

20 Plumley, J. A.; Ali-Torres, J.; Pohl, G.; Dannenberg, J. J. Capping Amyloid β-Sheets of the Tau-Amyloid Structure VQIVYK with Hexapeptides Designed To Arrest Growth. An ONIOM and Density Functional Theory Study. J. Phys. Chem. B 2014, 118, 3326–3334.

21 Zhu, L.; Song, Y.; Cheng, P. N.; Moore, J. S. Molecular Design for Dual Modulation Effect of Amyloid Protein Aggregation. J. Am. Chem. Soc. 2015, 137, 8062–8068.

22 Griner, S. L.; Seidler, P.; Bowler, J.; Murray, K. A.; Yang, T. P.; Sahay, S.; Sawaya, M. R.; Cascio, D.; Rodriguez, J. A.; Philipp, S.; Sosna, J.; Glabe, C. G.; Gonen, T.; Eisenberg, D. S. Structure-Based Inhibitors of Amyloid Beta Core Suggest a Common Interface with Tau. eLife 2019, 8, e46924.

23 Lu, J.; Cao, Q.; Wang, C.; Zheng, J.; Luo, F.; Xie, J.; Li, Y.; Ma, X.; He, L.; Eisenberg, D.; Nowick, J.; Jiang, L.; Li, D. Structure-Based Peptide Inhibitor Design of Amyloid-β Aggregation. Front. Mol. Neurosci. 2019, 12, 54.

24 Murray, K. A.; Hu, C. J.; Griner, S. L.; Pan, H.; Bowler, J. T.; Abskharon, R.; Rosenberg, G. M.; Cheng, X.; Seidler, P. M.; Eisenberg, D. S. De Novo Designed Protein Inhibitors of Amyloid Aggregation and Seeding. Proc. Natl. Acad. Sci. U. S. A. 2022, 119, e2206240119.

25 Taş, K.; Volta, B. D.; Lindner, C.; El Bounkari, O.; Hille, K.; Tian, Y.; Puig-Bosch, X.; Ballmann, M.; Hornung, S.; Ortner, M.; Prem, S.; Meier, L.; Rammes, G.; Haslbeck, M.; Weber, C.; Megens, R. T. A.; Bernhagen, J.; Kapurniotu, A. Designed Peptides as Nanomolar Cross-Amyloid Inhibitors Acting via Supramolecular Nanofiber Co-Assembly. Nat. Commun. 2022, 13, 5004.

26 Abskharon, R.; Pan, H.; Sawaya, M. R.; Seidler, P. M.; Olivares, E. J.; Chen, Y.; Murray, K. A.; Zhang, J.; Lantz, C.; Bentzel, M.; Boyer, D. R.; Cascio, D.; Nguyen, B. A.; Hou, K.; Cheng, X.; Pardon, E.; Williams, C. K.; Nana, A. L.; Vinters, H. V.; Spina, S.; Grinberg, L. T.; Seeley, W. W.; Steyaert, J.; Glabe, C. G.; Ogorzalek Loo, R. R.; Loo, J. A.; Eisenberg, D. S. Structurebased design of nanobodies that inhibit seeding of Alzheimer’s patient-extracted tau fibrils. Proc. Natl. Acad. Sci. U. S. A. 2023 120, e2300258120.

27 Reinke, A. A.; Gestwicki, J. E. Insight into amyloid structure using chemical probes. Chem. Biol. Drug. Des. 2011, 77, 399–411.

28 Shahnawaz, M.; Mukherjee, A.; Pritzkow, S.; Mendez, N.; Rabadia, P.; Liu, X.; Hu, B.; Schmeichel, A.; Singer, W.; Wu, G.; Tsai, A. L.; Shirani, H.; Nilsson, K. P. R.; Low, P. A.; Soto, C. Discriminating α-synuclein strains in Parkinson’s disease and multiple system atrophy. Nature 2020, 578, 273–277.

29 Yang, H.; Yuan, P.; Wu, Y.; Shi, M.; Caro, C. D.; Tengeiji, A.; Yamanoi, S.; Inoue, M.; DeGrado, W. F.; Condello, C. EMBER Multidimensional Spectral Microscopy Enables Quantitative Determination of Disease- and Cell-Specific Amyloid Strains. Proc. Natl. Acad. Sci. 2023, 120, e2300769120.

30 Li, B.; Ge, P.; Murray, K. A.; Sheth, P.; Zhang, M.; Nair, G.; Sawaya, M. R.; Shin, W. S.; Boyer, D. R.; Ye, S.; Eisenberg, D. S.; Zhou, Z. H.; Jiang, L. Cryo-EM of Full-Length α-Synuclein Reveals Fibril Polymorphs with a Common Structural Kernel. Nat. Commun. 2018, 9, 3609.

31 Boyer, D. R.; Li, B.; Sun, C.; Fan, W.; Zhou, K.; Hughes, M. P.; Sawaya, M. R.; Jiang, L.; Eisenberg, D. S. The α-synuclein hereditary mutation E46K unlocks a more stable, pathogenic fibril structure. Proc. Natl. Acad. Sci. U. S. A. 2020,117, 3592–3602.

32 Mitternacht, S. FreeSASA: An Open Source C Library for Solvent Accessible Surface Area Calculations. F1000Research 2016, 5, 189.

33 Zhou, J.; Grigoryan, G. Rapid Search for Tertiary Fragments Reveals Protein Sequence-Structure Relationships. Protein Sci. 2015, 24, 508–524.

34 Zhang, S. Q.; Kulp, D. W.; Schramm, C. A.; Mravic, M.; Samish, I.; DeGrado, W. F. The membrane-and soluble-protein helix-helix interactome: similar geometry via different interactions. Structure 2015, 23, 527–541.

35 Grigoryan, G.; Degrado, W. F. Probing designability via a generalized model of helical bundle geometry. J. Mol. Biol. 2011 405, 1079–1100.

36 Maguire, J. B.; Boyken, S. E.; Baker, D.; Kuhlman, B. Rapid Sampling of Hydrogen Bond Networks for Computational Protein Design. J. Chem. Theory Comput. 2018, 14, 2751–2760.

37 Leman, J. K.; Weitzner, B. D.; Lewis, S. M.; Adolf-Bryfogle, J.; Alam, N.; Alford, R. F.; Aprahamian, M.; Baker, D.; Barlow, K. A.; Barth, P.; Basanta, B.; Bender, B. J.; Blacklock, K.; Bonet, J.; Boyken, S. E.; Bradley, P.; Bystroff, C.; Conway, P.; Cooper, S.; Correia, B. E.; Coventry, B.; Das, R.; De Jong, R. M.; DiMaio, F.; Dsilva, L.; Dunbrack, R.; Ford, A. S.; Frenz, B.; Fu, D. Y.; Geniesse, C.; Goldschmidt, L.; Gowthaman, R.; Gray, J. J.; Gront, D.; Guffy, S.; Horowitz, S.; Huang, P.-S.; Huber, T.; Jacobs, T. M.; Jeliazkov, J. R.; Johnson, D. K.; Kappel, K.; Karanicolas, J.; Khakzad, H.; Khar, K. R.; Khare, S. D.; Khatib, F.; Khramushin, A.; King, I. C.; Kleffner, R.; Koepnick, B.; Kortemme, T.; Kuenze, G.; Kuhlman, B.; Kuroda, D.; Labonte, J. W.; Lai, J. K.; Lapidoth, G.; Leaver-Fay, A.; Lindert, S.; Linsky, T.; London, N.; Lubin, J. H.; Lyskov, S.; Maguire, J.; Malmström, L.; Marcos, E.; Marcu, O.; Marze, N. A.; Meiler, J.; Moretti, R.; Mulligan, V. K.; Nerli, S.; Norn, C.; Ó’Conchúir, S.; Ollikainen, N.; Ovchinnikov, S.; Pacella, M. S.; Pan, X.; Park, H.; Pavlovicz, R. E.; Pethe, M.; Pierce, B. G.; Pilla, K. B.; Raveh, B.; Renfrew, P. D.; Burman, S. S. R.; Rubenstein, A.; Sauer, M. F.; Scheck, A.; Schief, W.; Schueler-Furman, O.; Sedan, Y.; Sevy, A. M.; Sgourakis, N. G.; Shi, L.; Siegel, J. B.; Silva, D.-A.; Smith, S.; Song, Y.; Stein, A.; Szegedy, M.; Teets, F. D.; Thyme, S. B.; Wang, R. Y.-R.; Watkins, A.; Zimmerman, L.; Bonneau, R. Macromolecular Modeling and Design in Rosetta: Recent Methods and Frameworks. Nat. Methods 2020, 17, 665–680.

38 DiMaio, F.; Leaver-Fay, A.; Bradley, P.; Baker, D.; André, I. Modeling Symmetric Macromolecular Structures in Rosetta3. PLOS ONE 2011, 6, e20450.

39 Jullian, M.; Hernandez, A.; Maurras, A.;, Puget, K.; Amblard, M.; Martinez, J.; Subra, G. Nterminus FITC labeling of peptides on solid support: the truth behind the spacer. Tetrahedron Lett. 2009, 50,260–263.

40 Bousset, L.; Pieri, L.; Ruiz-Arlandis, G.; Gath, J.; Jensen, P. H.; Habenstein, B.; Madiona, K.; Olieric, V.; Böckmann, A.; Meier, B. H.; Melki, R. Structural and functional characterization of two alpha-synuclein strains. Nat. Commun. 2013, 4, 2575.

41 Yan, Y.; Zhang, D.; Zhou, P.; Li, B.; Huang, S. Y. HDOCK: a web server for protein-protein and protein-DNA/RNA docking based on a hybrid strategy. Nucleic Acids Res. 2017, 45, W365–W373.

42 Arosio P, Cukalevski R, Frohm B, Knowles TP, Linse S. Quantification of the concentration of Aβ42 propagons during the lag phase by an amyloid chain reaction assay. J. Am. Chem. Soc. 2014, 136, 219–225.

