## Supplementary material for "*De novo* Design of Peptides that Bind Specific Conformers of α-Synuclein": SI

### *De novo* Design of Peptides that Bind Specific Conformers of Amyloids; Methods and Application to α-Synuclein

**Table of Contents**

**Fig S1.** Interaction energies vs. packing scores of designed peptides to amyloid S1
**Fig S2.** Quadratic binding formula. S1

**Materials and Methods**
Preparation of αSyn monomer. S3
Preparation of fibrils. S3
Solid-phase peptide synthesis S3
Confocal microscopy imaging S3
Docking S4
Molecular dynamics simulation S4
Ligand depletion sedimentation assay S4
ThT fluorescence assay S5

**Supplementary Text** S6
Rosetta design scripts and settings

**References** S10

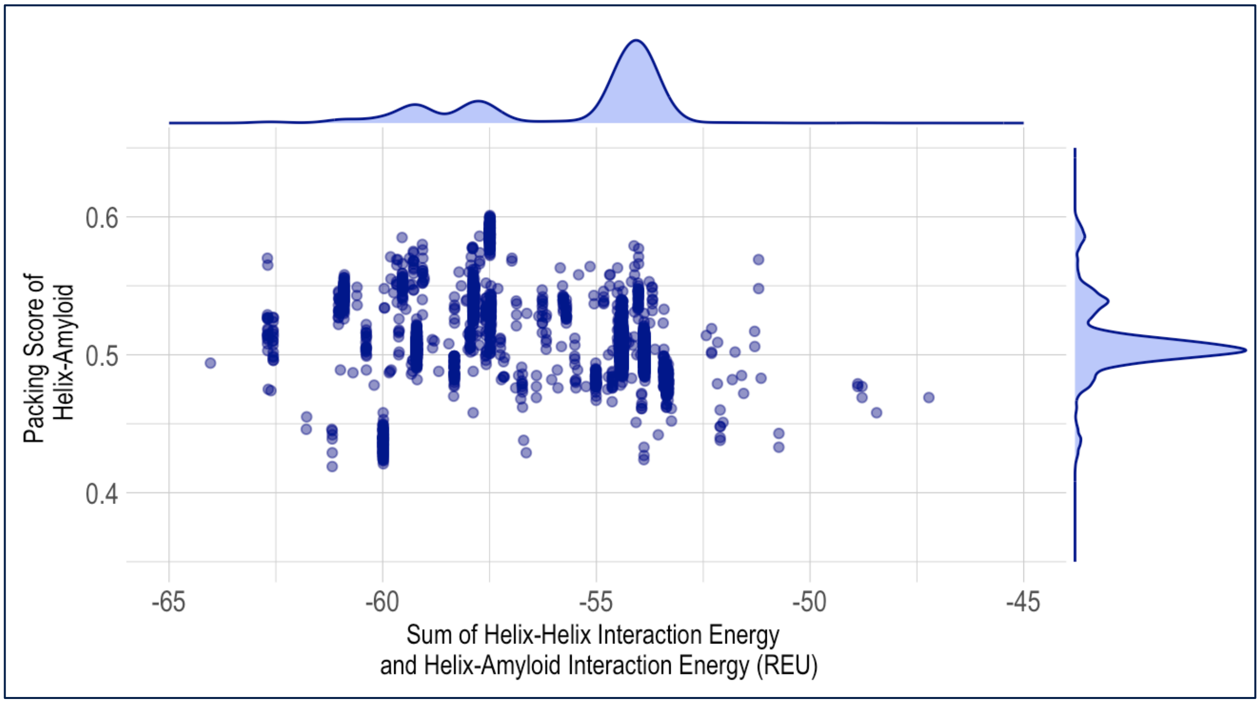


**Fig S1.** Interaction energies vs. packing scores of designed peptides to amyloid. Each point represents a different design model with plotted distributions on the x and y-axis. Each design model's interaction energy is plotted against the packing score.

$$\frac{[PL]}{[P]_T} = \frac{\left(K_D + \frac{[P]_T}{n} + [L]_T \pm \sqrt{\left(K_D + \frac{[P]_T}{n} + [L]_T\right)^2 - \frac{4[P]_T[L]_T]}{n}}\right)}{\frac{2[P]_T}{n}}$$

**Fig S2.** Quadratic binding formula to best fit ligand depletion sedimentation assay.

**Materials and Methods**

**Preparation of α-Syn monomers.**

α-Synuclein (α-Syn) fibrils were generated following established protocols to ensure consistent preparations.[1,2] In a concise description, the pET28a vector carrying the human α-synuclein gene was introduced into the BL21(DE3) *E. coli* strain, facilitating the overexpression of α-synuclein protein through IPTG induction. Subsequently, the cells were pelleted and suspended in an osmotic shock buffer (20 mM Tris-HCl, 40% sucrose, 2 mM EDTA at pH 7.2) and subjected to centrifugation. The resulting pellet was resuspended in cold water containing $MgCl_2$, followed by another round of centrifugation. The supernatant obtained was collected and subjected to lyophilization. E46K α-Syn was prepared the same way.

**Preparation of fibrils.**

*α-Synuclein:* The lyophilized α-synuclein monomers were reconstituted in a fibrillization solution containing 15 mM tetrabutylphosphonium bromide, resulting in a final concentration of 300 μM α-synuclein. Over a period of seven days, fibrils were grown through continuous shaking at room temperature in non-stick Eppendorf tubes.

*Aβ40, 0N3R tau, α-Synuclein (RIBBON), α-Synuclein (FIBRIL):* Previously reported procedures were followed.[3,4,5]

**Solid-phase peptide synthesis.**

The designed peptides were synthesized using Fmoc solid-phase peptide synthesis. An automated peptide synthesizer (Biotage Initiator+ Alstra peptide synthesizer) was employed, utilizing TentagelS RAM resin at a 0.05 mmol scale (0.10 g with 0.3 mmol/g loading size). Each amino acid was coupled in five equiv Fmoc-protected amino acid with five equiv of HCTU and ten equiv of *N,N*-diisopropylethylamine (DIPEA) in DMF relative to the peptide functional sites, at 75 ºC for 5 min. Deprotection was achieved by treating the resin-bound peptides with 4.5 mL of 20% 4-methylpiperidine in DMF at 70 ºC for 5 min.

For FITC-tagged peptides, Fmoc-beta-alanine-OH was coupled and Fmoc-deprotected, then the N-terminus was conjugated with 2.5 equiv FITC along with 5 equiv of DIPEA relative to the peptide functional site for 5 h at room temperature on resin. The resin was washed with three rinses of DMF and four rinses of $CH_2Cl_2$ before being dried for 1 h.

Peptide cleavage was performed following a standard protocol in a solution of 95% trifluoroacetic acid (TFA), 2.5% $H_2O$, 2.5% triisopropylsilane (TIPS), and 20 mg/mL dithiothreitol (DTT) for 1 h at room temperature. The crude peptide was then precipitated in cold diethyl ether and subsequently lyophilized. The resulting crude peptide was dissolved in $H_2O$ + 0.1% TFA and purified by preparative reverse-phase HPLC using a Vydac C4 column (22 mm × 250 mm, 10 μm particle size) and a linear gradient of 5-100% MeCN + 0.1% TFA (with the remaining fraction $H_2O$ + 0.1% TFA) over 25 min at a flow rate of 10 mL/min. The peptide eluted at approximately 40% MeCN. Fractions containing the peptide were combined and peptide purity was assessed by MALDI mass spectrometry and analytical HPLC. The combined fractions were frozen in liquid nitrogen followed by lyophilization. Approximately 20 mg of purified peptide was obtained as a white powder from the 0.05 mmol synthesis scale, corresponding to an overall yield of approximately 5%.

**Confocal microscopy imaging.**

Fluorophore-conjugated peptides were prepared in 1x PBS varying concentrations (128, 64, 32, 16, 8, 4, 2, 1, 0.1, 0.01 nM). For the experimental setup, 50-μL aliquots of each peptide solution were transferred into each wells of a 384-well plate (Corning™ BioCoat™ 384-Well, Collagen Type I-Treated, Flat-Bottom Microplate). Subsequently, 1-μL fibril solution was added to each well, achieving a final fibril concentration of 600 nM. Thorough mixing of the fibril and peptide solutions was achieved by pipetting up and down. The plate was centrifuged at 50 x g to pellet the fibrils.

Imaging of the fibril-peptide complexes was performed using a Leica SP8 confocal microscope equipped with a 40× water immersion lens (1.1 NA), white light and 405 nm lasers, and a HyD detector at 512 x 512 with a 1x zoom factor and a LightGate (0.5 to 18 ns). Excitation and emission detection wavelengths were collected at 490/525 nm.

**Docking**
*MOE:* Rigid-body protein-protein docking was performed by MOE software 2022.02 (Chemical Computing Group ULC, Montreal, Quebec, Canada). The receptor structure was obtained from alpha-syn fibril structure (PDB ID: 6CU7) and the ligand structure was imported from the designed trimeric helical bundle. Refinement was performed using rigid body and the top 100 poses were visually examined.
*HDOCK:* Molecular docking simulations were conducted using HDOCK.[6] The simulations were run on default parameters for protein-protein docking and the top 100 predictions were generated. The poses corresponding to the lowest energy scores is shown in the figure.

**Molecular dynamics simulations.**
Four copies of each designed peptide were simulated in complex with ten fibrils (except peptide 5, which was simulated with 9 fibrils because the peptide is much shorter in length and does not cross over multiple fibrils). The *N*- and *C*-termini of the amyloid regions were capped with *N*-methyl and acetyl group, respectively. The *N*-termini of the designed peptides were capped with acetyl groups, while their *C*-termini were amidated.
The MD system was prepared using tleap.[7,8] The simulation box was built by solvating each complex with OPC model waters[9] with 8 Å padding from the protein, and the system was charge-neutralized. All simulations were performed in Amber18 with the ff19SB forcefield.[10] Simulations began with 1,000 restrained steepest-descent minimization steps before switching to a maximum of 7,000 steps in conjugate gradient minimization. The system was then heated up from 150K to 300 K over 50 ps in the NVT ensemble with Langevin thermostat control of temperature, using a 1 fs integration timestep. The system was then switched to the NPT ensemble, and pressure was maintained at 1 atm using the Monte Carlo barostat. Peptide residues were restrained with a 10 kcal/(mol·Å2) force constant initially and very slowly ramped down to 0 kcal/(mol·Å2) over 6 equilibration steps, comprising 3ns in total. Amyloid regions were restrained with a 10 kcal/(mol·Å2) force constant throughout the whole simulation to maintain their positional integrity.
Following minimization and equilibration, the simulation was carried out for a 500 ns production run under periodic boundary conditions with 2 fs timesteps. The SHAKE algorithm[11] was used to restrain hydrogens, the Particle Mesh Ewald method[12,13,14,15] was used to calculate long-range electrostatics, and non-bonded interactions were cut off at 10 Å. Three independent simulations were performed for each design.

**Ligand depletion sedimentation assay**
The WT αSyn fibril was diluted 2-fold with a PBS solution containing 1 μM peptide 4*, resulting in concentrations of 100, 50, 25, 12.5, 6.25, 3.125, 1.5625, and 0.78 μM. After incubating the mixture at room temperature for 30 minutes, it was centrifuged at 14,000 rpm for 20 minutes to pellet the fibrils and fibril-bound peptide 4*. The supernatant, containing the fibril unbound peptides, was carefully collected. The fluorescence signal associated with the FITC-labeled peptide in the supernatant was quantified by measuring the absorbance of FITC at 490 nm. Each experiment was conducted in duplicates, and the same procedure was followed for the scramble* peptide.

**ThT fluorescence assay.**

α-Synuclein fibrillization was monitored using the FLUOstar OMEGA microplate reader. A 50 μL of 300 μM α-synuclein monomers containing 15 μM Thioflavin T (ThT) was transferred to a 384-well plate. The ThT fluorescence resulting from 440 nm excitation and 490 nm emission was recorded at every 10-min interval over the course of 6000 min period. The plate was agitated for 5 min between each data collections.

**Supplementary Text**

*RosettaScript (design_hbnet.xml)*

```
<ROSETTASCRIPTS>
	<SCOREFXNS>
		<ScoreFunction name="ref15" weights="ref2015" symmetric="1">
		</ScoreFunction>
	</SCOREFXNS>

	<RESIDUE_SELECTORS>
	</RESIDUE_SELECTORS>

	<TASKOPERATIONS>
		<InitializeFromCommandline name="ifcl"/>
		<ReadResfile name="resfile" filename="resfile"/>
		<ExtraRotamersGeneric name="extrachi" ex1="1" ex2="1" ex1_sample_level="1"
ex2_sample_level="1" extrachi_cutoff="14"/>
		<IncludeCurrent name="include_curr" />
	</TASKOPERATIONS>

	<FILTERS>
		<PackStat name="pstat" threshold="0.30" repeats="10" /> Common wisdom says
0.65 is a good number to look out for.
		<PackStat name="pstat_mc" threshold="0" repeats="10"/>
		<NetCharge name="net_charge" confidence="0"/>
		<ScoreType name="total_score" scorefxn="ref15" score_type="total_score"
threshold="300"/>
		<ScoreType name="total_score_1" scorefxn="ref15" score_type="total_score"
				threshold="0"/>
	</FILTERS>

	<MOVERS>
		<SetupForSymmetry name="setup_symm" definition="query2_end.symm"/>
		<HBNetStapleInterface scorefxn="ref15" name="hbnet_interf" monte_carlo="true"
hb_threshold="-0.5" write_network_pdbs="false" write_cst_files="false"
			design_residues="NSTQHYWKRDE" min_networks_per_pose="1"
max_networks_per_pose="5" use_aa_dependent_weights="true"
			min_core_res="1" min_network_size="3" max_unsat_Hpol="1"
onebody_hb_threshold="-0.3" task_operations="ifcl,resfile,include_curr" />
		<MultiplePoseMover name="MPM_design" max_input_poses="100">

			<ROSETTASCRIPTS>
				<SCOREFXNS>
					<ScoreFunction name="ref15" weights="ref2015">
					</ScoreFunction>
				</SCOREFXNS>

				<RESIDUE_SELECTORS>
				</RESIDUE_SELECTORS>

				<TASKOPERATIONS>
					<InitializeFromCommandline name="ifcl"/>
```

```
<ReadResfile name="resfile" filename="resfile"/>
<ExtraRotamersGeneric name="extrachi" ex1="1" ex2="1"
ex1_sample_level="1" ex2_sample_level="1" extrachi_cutoff="14"/>
<IncludeCurrent name="include_curr" />
</TASKOPERATIONS>

<FILTERS>
<PackStat name="pstat" threshold="0.30" repeats="10" />
Common wisdom says 0.65 is a good number to look out for.
<PackStat name="pstat_mc" threshold="0" repeats="10"/>
<NetCharge name="net_charge" confidence="0"/>
<ScoreType name="total_score" scorefxn="ref15"
score_type="total_score" threshold="300"/>
<ScoreType name="total_score_1" scorefxn="ref15"
score_type="total_score"
threshold="0"/>
</FILTERS>

<MOVERS>
<SetupForSymmetry name="setup_symm"
definition="query2_end.symm"/>
<PackRotamersMover name="pack" scorefxn="ref15"

task_operations="ifcl,resfile,include_curr,extrachi"/>
<PackRotamersMover name="pack_fast" scorefxn="ref15"

task_operations="ifcl,resfile,include_curr"/>
<MinMover name="min_bb" scorefxn="ref15"
tolerance="0.0000001" max_iter="1000" chi="false" bb="true">
<MoveMap name="map_bb">
<Span begin="1" end="120" bb="false"
chi="false" />
<Span begin="121" end="999" bb="true"
chi="false"/>
<Jump number="2" setting="1" />
</MoveMap>
</MinMover>
<Idealize name="idealize"/>
<MinMover name="min_sc" scorefxn="ref15"
tolerance="0.0000001" max_iter="1000" chi="true" bb="false">
<MoveMap name="map_sc">
<Span begin="1" end="120" bb="false"
chi="false" />
<Span begin="121" end="999" bb="false"
chi="true"/>
</MoveMap>
</MinMover>
<MinMover name="min_sc_bb" scorefxn="ref15"
tolerance="0.0000001" max_iter="1000" chi="true" bb="true">
<MoveMap name="map_sc_bb">
```

```
                    <Span begin="1" end="120" bb="false"
chi="false" />
                    <Span begin="121" end="999" bb="true"
chi="true"/>
                    <Jump number="2" setting="1" />
                </MoveMap>
            </MinMover>
            <ParsedProtocol name="parsed_pack_fast" >
                <Add mover_name="pack_fast"/>
                <Add mover_name="min_bb"/>
            </ParsedProtocol>
            <ParsedProtocol name="parsed_pack" >
                <Add mover_name="pack"/>
                <Add mover_name="min_bb"/>
                <Add mover_name="min_sc"/>
            </ParsedProtocol>
            <GenericMonteCarlo name="pack_mc" preapply="0"
trials="3" temperature="0.03"

    filter_name="pstat_mc" sample_type="high" mover_name="parsed_pack">
                    <Filters>
                        <AND filter_name="total_score_1"
temperature="15" sample_type="low"/>
                    </Filters>
            </GenericMonteCarlo>
            <GenericMonteCarlo name="pack_fast_mc" preapply="0"
trials="2" temperature="0.03"

    filter_name="pstat_mc" sample_type="high" mover_name="parsed_pack_fast">
                    <Filters>
                        <AND filter_name="total_score_1"
temperature="15" sample_type="low"/>
                    </Filters>
            </GenericMonteCarlo>

        </MOVERS>
        <APPLY_TO_POSE>
        </APPLY_TO_POSE>

        <PROTOCOLS>
            <Add mover="setup_symm"/>
            <Add mover_name="parsed_pack_fast"/>
            <Add mover_name="pack_fast_mc"/>
            <Add mover_name="pack_mc"/>
            <Add mover_name="min_sc_bb"/>
            <Add filter_name="pstat"/>
            <Add filter_name="net_charge"/>
            <Add filter_name="total_score"/>
        </PROTOCOLS>

        <OUTPUT scorefxn="ref15"/>
```

```
                    </ROSETTASCRIPTS>
                </MultiplePoseMover>
        </MOVERS>
        <APPLY_TO_POSE>
        </APPLY_TO_POSE>

        <PROTOCOLS>
                <Add mover="setup_symm"/>
                <Add mover_name="hbnet_interf"/>
                <Add mover_name="MPM_design"/>
        </PROTOCOLS>

        <OUTPUT scorefxn="ref15"/>
</ROSETTASCRIPTS>
```

*Resfile*

```
NATRO
start

* C ALLAAxc
# this line means that ALL (*) residues on chain C can be designed
# with ALL AA (amino acids) except cysteine (XC) -- to avoid disulfide linkages


# the following lines are commented out, but this is what the rest of the
# resfile would look like if you wanted to design specific amino acids on to
# specific resdiue locations

#1 B PIKAA KREQS
#2 B PIKAA ALKRQEMWFS
#4 B PIKAA ALKRQEMWFS
#5 B PIKAA KREQS
#6 B PIKAA ALKRQEMWFS
#8 B PIKAA KREQS
#9 B PIKAA KREQS
#12 B PIKAA KREQS
#13 B PIKAA ALKRQEMWFS
#15 B PIKAA ALKRQEMWFS
#16 B PIKAA KREQS
```

*Symmetry*

```
symmetry_name query2_sym_pseudo3fold
E = 2*VRT_0_base + 1*(VRT_0_base:VRT_1_base) + 1*(VRT_0_base:VRT_2_base)
anchor_residue COM
virtual_coordinates_start
xyz     VRT_0       1.000000,0.000000,0.000000      0.000000,1.000000,0.000000
52.136375,41.582250,25.056844
xyz     VRT_0_base  1.000000,0.000000,0.000000      0.000000,1.000000,0.000000
52.136375,41.582250,25.056844
xyz     VRT_1       0.999493,0.031702,-0.002818     -0.031683,0.999477,0.006443
51.091563,40.926625,15.449813
```

```
xyz	VRT_1_base	0.999493,0.031702,-0.002818	-0.031683,0.999477,0.006443
51.091563,40.926625,15.449813
xyz	VRT_2	0.999493,-0.031693,0.003027	0.031712,0.999477,-0.006333
53.174281,42.266344,34.662687
xyz	VRT_2_base	0.999493,-0.031693,0.003027	0.031712,0.999477,-0.006333
53.174281,42.266344,34.662687
virtual_coordinates_stop
connect_virtual JUMP_0_to_subunit VRT_0_base SUBUNIT
connect_virtual JUMP_1_to_subunit VRT_1_base SUBUNIT
connect_virtual JUMP_2_to_subunit VRT_2_base SUBUNIT
connect_virtual JUMP_0_to_com VRT_0 VRT_0_base
connect_virtual JUMP_1_to_com VRT_1 VRT_1_base
connect_virtual JUMP_2_to_com VRT_2 VRT_2_base
connect_virtual JUMP_1 VRT_0 VRT_1
connect_virtual JUMP_2 VRT_0 VRT_2
set_dof JUMP_0_to_com x y z
set_dof JUMP_0_to_subunit angle_x angle_y angle_z
set_jump_group JUMPGROUP1  JUMP_0_to_subunit JUMP_1_to_subunit JUMP_2_to_subunit
set_jump_group JUMPGROUP2  JUMP_0_to_com JUMP_1_to_com JUMP_2_to_com
```

#### References


1 Huang, C.; Ren, G.; Zhou, H.; Wang, C. C. A new method for purification of recombinant human α-synuclein in Escherichia coli. *Protein Expr. Purif.* **2005**, *42*, 173–177.

2 Kloepper, K. D.; Woods, W. S.; Winter, K. A.; George, J. M.; Rienstra, C. M. Preparation of α-synuclein fibrils for solid-state NMR: expression, purification, and incubation of wild-type and mutant forms. *Protein Expr. Purif.* **2006**, *48*, 112–117.

3 Wang, T.; Jo, H.; DeGrado, W. F.; Hong, M. Water Distribution, Dynamics and Interactions with Alzheimer's βAmyloid Fibrils Investigated by Solid-State NMR. *J. Am. Soc. Chem.* **2017**, *139*, 6242–6252.

4 Dregni, A. J.; Wang, H. K.; Wu, H.; Duan, P.; Jin, J.; DeGrado, W. F.; Hong, M. Inclusion of the C-Terminal Domain in the β-Sheet Core of Heparin-Fibrillized Three-Repeat Tau Protein Revealed by Solid-State Nuclear Magnetic Resonance Spectroscopy. *J. Am. Chem. Soc.* **2021**, *143*, 7839–7851.

5 Li, B.; Ge, P.; Murray, K. A.; Sheth, P.; Zhang, M.; Nair, G.; Sawaya, M. R.; Shin, W. S.; Boyer, D. R.; Ye, S.; Eisenberg, D. S.; Zhou, Z. H.; Jiang, L. Cryo-EM of Full-Length α-Synuclein Reveals Fibril Polymorphs with a Common Structural Kernel. *Nat. Commun.* **2018**, *9*, 3609.

6 Yan, Y.; Zhang, D.; Zhou, P.; Li, B.; Huang, S. Y. HDOCK: a web server for protein-protein and protein-DNA/RNA docking based on a hybrid strategy. *Nucleic Acids Res.* **2017**, *45*, W365–W373.

7 D.A. Case, I.Y. Ben-Shalom, S.R. Brozell, D.S. Cerutti, T.E. Cheatham, III, V.W.D. Cruzeiro, T.A. Darden, R.E. Duke, D. Ghoreishi, M.K. Gilson, H. Gohlke, A.W. Goetz, D. Greene, R Harris, N. Homeyer, Y. Huang, S. Izadi, A. Kovalenko, T. Kurtzman, T.S. Lee, S. LeGrand, P. Li, C. Lin, J. Liu, T. Luchko, R. Luo, D.J. Mermelstein, K.M. Merz, Y. Miao, G. Monard, C. Nguyen, H. Nguyen, I. Omelyan, A. Onufriev, F. Pan, R. Qi, D.R. Roe, A. Roitberg, C. Sagui, S. Schott-Verdugo, J. Shen, C.L. Simmerling, J. Smith, R. SalomonFerrer, J. Swails, R.C. Walker, J. Wang, H. Wei, R.M. Wolf, X. Wu, L. Xiao, D.M. York and P.A. Kollman (2018), AMBER 2018, University of California, San Francisco.

8 Salomon-Ferrer, R.; Case, D. A.; Walker, R. C. An overview of the Amber biomolecular simulation package. *Wires Comput Mol. Sci.* **2013**, *3*, 198–210.

9 Izadi, S.; Anandakrishnan, R.; Onufriev, A.V. Building Water Models: A Different Approach. *J. Phys. Chem. Lett.* **2014**, *5*, 3863–3871.

10 Tian, C.; Kasavajhala, K.; Belfon, K.; Raguette, L.; Huang H.; Migues, A.; Bickel, J.; Wang, Y.; Pincay, J.; Wu, Q.; Simmerling, C. ff19SB: Amino-Acid-Specific Protein Backbone Parameters Trained against Quantum Mechanics Energy Surfaces in Solution. *J. Chem. Theory Comput.* **2020**, *16*, 528–552.

11 Ryckaert, J.-P.; Ciccotti, G.; Berendsen, H.J.C. Numerical integration of the cartesian equations of motion of a system with constraints: Molecular dynamics of n-alkanes. *J. Comput. Phys.* **1977**, *23*, 327–341.

12 Darden, T.; York, D.; Pedersen, L. Particle mesh Ewald–an Nlog(N) method for Ewald sums in large systems. *J. Chem. Phys.* **1993**, *98*, 10089–10092.

13 Essmann, U.; Perera, L.; Berkowitz, M.L.; Darden, T.; Lee, H.; Pedersen, L.G. A smooth particle mesh Ewald method. *J. Chem. Phys.* **1995**, *103*, 8577–8593.

14 Crowley, M.F.; Darden, T.A.; Cheatham, T.E. III; Deerfield, D.W. II. Adventures in improving the scaling and accuracy of a parallel molecular dynamics program. *J. Supercomput.* **1997**, *11*, 255–278.

15 Sagui, C.; Darden, T.A. in Simulation and Theory of Electrostatic Interactions in Solution, Pratt, L.R.; Hummer, G., Eds., pp 104–113. American Institute of Physics, Melville, NY, 1999.
